# Relationship between depressive symptoms and cumulative 24-hour urinary norepinephrine excretion level among undergraduate medical students in Uganda

**DOI:** 10.1101/695825

**Authors:** Mutiat O Owoola-Ajirotutu, Alfred O Okpanachi, Akeem G Owoola, Godfrey Z Rukundo, Sadiq Yusuf

**Author notes:** Corresponding author (SY). Women’s Hospital International & Fertility Centre, Bukoto-Kisasi Road, P.O Box 16233, Bukoto-Kampala, Uganda. These authors contributed equally to this work (Mutiat O Owoola-Ajirotutu, Sadiq Yusuf). These authors also contributed equally to this work (Alfred O Okpanachi, Akeem G Owoola, Godfrey Z Rukundo).

## Abstract

**Background:** Depression is a serious mental health problem in different parts of the world and has been reported to be rising among undergraduate medical students. The incidence of depression has not only been linked to psychosocial factors but also to biological factors, such as altered urinary levels of norepinephrine. This study was carried out to determine the prevalence of depression among undergraduate medical students in Uganda and examine the relationship between depressive symptoms and 24-hour urinary norepinephrine excretion levels in the participants.

**Methods:** One hundred and sixteen undergraduate medical students (75 males and 41 females) of Kampala International University, in southwestern Uganda were evaluated for depression using the 21-item Beck Depression Inventory-II (BDI) questionnaire. Twenty-four-hour urine collections from each participant were assayed for norepinephrine excretion levels. Descriptive statistics and Pearson correlation coefficient were computed to examine the data obtained.

**Results:** The results of this study showed that, a total of 33 participants (28.4%) have depressive symptoms. Students with depressive symptoms had higher but not significant 24-hour urinary mean norepinephrine excretion levels than those without depressive symptoms (121.97±51.48μg/day Vs 87.58±18.64 μg/day, P>0.05). There was a positive weak relationship between BDI scores and 24-hour urinary norepinephrine levels (r= 0.21, p = 0.28). Regression models accounting for socio-demographic characteristics indicated that, type of accommodation, marital status, relationship with parents, educational sponsorship may be risk factors for depressive symptoms observed in the participants.

**Conclusions:** These results suggest that increased urinary norepinephrine excretion and other psychosocial factors may be associated with depressive symptoms. Measurements of 24-hour urinary norepinephrine excretion may serve as an integrative parameter in diagnosing and management of patients with depression.

## Introduction

Depression is one of the most common mental disorders affecting people of different ages, gender and socio-cultural settings in both developed and developing countries. It has been declared by the World Health Organization as one of the global leading causes of morbidity and mortality contributing significant economic burden on societies worldwide [1–5]. Studies have suggested that medical students are at high risk of developing depression and this has been associated with significant reduction in productivity, disruption in learning, poor interpersonal relationships with peers and consequently poor academic grades and in some cases termination of schooling which often leads to substance abuse and suicidal behavior [6–8]. Since depression among medical students have such an obvious negative effect on function in medical school and later in clinical practice, it is important to examine the prevalence, causes and diagnosis of the disorder in medical students. The prevalence and risk factors for depression in Ugandan university students has not been well studied in Uganda. The few available studies have examined depression amongst adolescents in secondary schools [9], traumatized individuals [10] and among persons living with human immune deficiency virus (HIV) infection [11]. Although psychosocial problems had been reported among the undergraduate students in Uganda [12], no study has examined the link between socio-demographic factors and biological effects of the disease in this population.

Previous research has suggested that enhanced activity of the hypothalamic–pituitary–adrenal (HPA) axis with concomitant increased concentrations of catecholamines such as serotonin (5-HT), norepinephrine (NE), and dopamine (DA) plays an etiological role in the onset and development of depressive symptoms [13, 14]. Serotonin and DA have been the most studied neurotransmitters in depression, however, converging lines of evidence suggest that the NE pathway is of major importance in the pathophysiology and treatment of depressive disorder [15–18]. The noradrenergic system uses NE as the main chemical messenger and serves multiple brain functions including arousal, motivation, attention, mood, learning, memory and stress response [19]. Higher NE levels have been linked to low socioeconomic status [20]. Neuroactivities of NE-selective tricyclic antidepressants such as desipramine and nortriptyline indicates that NE could be majorly involved in NE neurotransmission in depression [21]. These studies provide a rationale for measuring NE output in depression and invoke NE as player in etiology of this disorder.

Studies have highlighted a need for biomarkers in psychiatry to enhance patient management and ensure treatment success [22, 23]. While urinary measures of neurotransmitters are not a direct assessment of central nervous system activities, urinary excretion of neurotransmitters or their metabolites have been characterized as biomarkers of various neurological conditions [24, 25]. Thus, the urinary excretion of NE may act as a biomarker in diagnosing and management of patients with depression. Given the high prevalence and costs of depression, the impairment associated with depression, and the difficulty in treating depression once it has developed, efforts to improve the early detection of depression and treat it as soon as possible are warranted. This study was therefore, designed to examine the association between prevalence of depressive symptoms and 24-hour (24-h) urinary NE excretion levels in a representative sample of undergraduate medical students in Uganda. The results obtained maybe important in the efforts to identify the causes and diagnosis of the disorder among medical students.

## Materials and methods

### Participants

One hundred and sixteen (116) male and female students were studied in the Faculty of Biomedical Sciences in the School of Health Sciences of Kampala International University, Uganda. The students in biomedical sciences are made up of three group of classes which include students in semesters one to three respectively. Each class was considered to be a cluster and using the appropriate allocation method of sampling, one class was randomly selected from the three classes. Thus, all the 169 students in the selected class constituted the participants of the present study. Of these, the relationship of depressive symptoms with 24-h urinary NE excretion levels was evaluated in 75 males and 41 females (n = 116). These participants returned the questionnaires and their urine samples were verified to meet the collection and storage criteria during the 24-h collection period. All participants had no history of cardiovascular or metabolic diseases.

A coded self-administered questionnaire requesting information on the participant’s age, sex, marital status, family type, financial support, parental loss, occupation, type of residence and nationality were filled by all participants. To ensure reliability in the participant’s response, validation and reliability studies were done in pre-test on 25 students in another class. Cronbach’s alpha was used to measure the reliability of the questions. The results showed that the questionnaire has high validation and reliability scores with a Cronbach’s alpha coefficient of 0.8. All experimental protocols were approved by the Institutional Research Ethics Committee (IREC) of Mbarara University of Science and Technology, Uganda. Informed consent was obtained from all participants prior to participation in this study. Confidentiality of information was maintained by ensuring that codes but not names were used to label the study tools.

### Determination of depressive symptoms

The prevalence of depressive symptoms among participants was determined using the Beck Depression Inventory (BDI) scale-II. The BDI scale is a well-validated and widely used 21-item self-report instrument developed to evaluate the presence and severity of depression in adults and adolescents in non-clinical settings [26–30]. In this study, participants were advised to choose an option for each question that best described their feelings over the preceding week. Responses to the 21 items were summed up to give their depressive status on a 4-point scale giving a maximum score of 63. Participants with BDI score of 0-9 were categorized as normal, scores of 10-18 were indicative of mild mood disturbance, scores of 19-29 were indicative of moderate depression, scores of 30-40 were indicative of severe depression and those with score of 40 and over were categorized as having very severe depression.

The prevalence of suicide ideation was measured with a frequency count of “yes” response to each of the following questions: a) Have you ever experienced suicide thoughts along with the wish to end your life by suicide? b) Did you experience suicide thoughts along with the wish to end your life by suicide last week? The participants found to have depressive symptoms or suicide ideation were instructed to discuss these symptoms with a physician for further evaluation and treatment.

### 24-hour urine collection

Participants were provided with 3-liter opaque collection jugs containing a preservative (10 ml of 6N hydrochloric acid, pH < 3.5) and a smaller 100 ml specimen bottles containing no preservative. The 3-litre collection jug was labelled with a code similar to that of the BDI and questionnaire earlier completed by the participant. To ensure urine collection during a typical day under typical conditions, the participants were asked to collect urine in their residential environment during a 24–h period.

They were instructed to collect all urine passed during the target period (Saturday to Sunday when there are no scheduled lectures or university activities). The collection of the 24-h urine started with the participant voiding (completely emptying bladder) and discarding the first urine passed on Saturday morning. Thereafter, they were instructed to collect all of the urine passed during that day and night including urine passed during bowel movements, up to and including the first voiding of the following day (Sunday). Each urine sample was first collected into the smaller container for ease of convenience and immediately transferred into the 3-litre collection jug containing preservative. The participants were asked to keep the urine collection jug tightly closed and refrigerated or kept in a cool place throughout the collection period. No dietary restrictions were enforced; however, participants were advised to discontinue taking all medications for an interval of at least 12 hours preceding the urine collection period.

To ensure compliance with the collection protocol, mobile phone numbers of every five participants were assigned to a research assistant who sent four reminder messages at different times within the 24-h period to continue with collection and to verifying that the urine jugs were properly stored. If it was discovered that, a participant did not adhere to the collection protocol or if the volume collected was less than 1 litre, the participant was asked to repeat the collection the following weekend. In the laboratory, the volume of cumulative 24-h urine samples was recorded and stirred thoroughly, separated into 10 ml aliquots in a sterile sample bottles, and then stored in the laboratory refrigerator prior to analysis.

### Measurement of 24-hour urinary norepinephrine

Quantitative measurement of NE in each urine sample was performed by enzyme immunoassay (EIA) following a protocol described in detail by the manufacturer of the assay kit (Abnova, UK). Westermann, et al [25] established the accuracy and reproducibility of the enzyme linked immunoassay methodology for NE as compared to previously validated high-pressure liquid chromatography (HPLC) methodology. The authors concluded that EIA measures for urinary NE are appropriate for clinical applications as they were rapid, accurate, and reproducible.

#### Sample pre-treatment

An aliquot (10 μl) of working standard and 10 μl of urine samples were added to borate-coated wells of a macrotiter plate. To these, 250 μl of double distilled deionised water was added. 50μl of assay and extraction buffer (0.1 mol/L Tris-HCL buffer, 0.7 mol/L NaCl, 0.1 mol/L EDTA, 0.3 mmol/L Na_2_S_2_O_5_, pH 9.3) were added to each well of the plate respectively. The plates were covered with adhesive foil and incubated for 30 min at room temperature on a shaker (approx. 600 rpm) after which the plates were emptied and blotted dry by taping the inverted plate on absorbent material. Thereafter, 150 μl of acylation buffer was added to the wells respectively. The plates were incubated for further 15 min at room temperature on a shaker (approx. 600 rpm). Wash buffer (1 mL) was pipetted into the wells and the plates incubated for 10 min at room temperature on a shaker (approx. 600 rpm). 150 μl of hydrochloric acid was added into the wells and the plates were covered with adhesive foil. The plates were further incubated for 10 min at room temperature on a shaker (approx. 600 rpm) after which the foil was discarded and 20 μl of the supernatant was removed for subsequent norepinephrine EIA.

#### Enzyme Immunoassay

The enzyme solution (25 μl) was added into all wells of the NE microtiter strips followed by the addition of 20 μl of the extracted standards and urine samples into the appropriate wells. The solution was incubated at 37 °C for 30 min on a shaker. NE antiserum (50 μl) was added to the wells and covered with adhesive foil. The preparation was further incubated for 2 hours at room temperature on a shaker (approx. 600 rpm). Thereafter, the foil was removed and the content of the wells aspirated. The plates were then washed 3x by adding 300 μl of wash buffer (0.02 mol/L Tris-HCl buffer, 0.1 mol/L NaCl, 5 mmol/L KCl, 0.2% Tween 80, pH 7.3) discarding the content and blotting dry each time by tapping the inverted plate on absorbent material. 100 μl of the enzyme conjugate were then added to all the wells. The plates were incubated for 30 min at room temperature on a shaker. The contents of the wells were discarded and the plates were washed 3x by adding 300 μl of wash buffer, discarding the content and blotting dry each time by tapping the inverted plate on absorbent material. 100 μL of the substrate was added to all wells and incubated for 25 ± 5 min at room temperature on a shaker. The reaction was stopped when an orange color was developed in positive wells by adding 100 μL of the stop solution to each well.

The microtiter plate was shaken to ensure a homogeneous distribution of the solution. The developed color intensity is proportional to the NE concentration in the sample. The absorbance of the solution in the wells were read within 10 minutes, using a microplate reader set to 540 nm. Quantification of unknown samples was achieved by comparing their absorbance with a standard curve prepared with known standard concentrations. The standard curve was obtained by plotting the absorbance readings against the corresponding standard concentrations. The concentrations of NE in the urine samples were read directly from the standard curve. The normal reference range for the assay was < 90 μg/day for norepinephrine.

### Statistical analysis

Analysis of data was performed with Microsoft Excel for Mac (2016). The Data obtained for socio-demographic characteristics of participants, 24-h NE levels and prevalence of depressive symptoms were expressed as mean ± standard deviation (SD) or number or percentage. Differences in the amount of NE in 24-h urine samples in normal and individuals exhibiting depressive symptoms were analysed using independent t-test. Correlations between depressive symptoms and levels of NE in 24-h urine samples were assessed using Pearson correlation coefficient and Chi-square tests to identify significant predictors of depression. Odds ratios (OR) and 95% confidence intervals were also calculated for each variable. All p-values were two-tailed, with a p-value of < 0.05 considered to be of statistical significance.

## Results

The study group included 75 men and 41 women representing 64.7% and 35.3% respectively. The age of participants ranged from 17 to 45 years with a mean (±SD) of 24.6±5.7 years. The socio-demographic data of all 116 participants are shown in Table 1. A total of 20 (17.2%) of the participants are married, 52 (44.8%) are single, 42 (36.2%) are in a relationship and 2 (1.7%) are divorced. Of all the participants, 81 (69.9%) were from a polygamous family and 35 (30.1%) were from a monogamous family. Majority of the participants were Ugandans (78.5%) followed by Kenyans and Nigerians (10%) respectively. They mainly lived in rented-room or house outside the campus (81.9%) while 10.3% and 7.8% were living in the university hostel and with parents or relatives respectively. Among participants that have religion, 80.2% and 16.4% of participants consider themselves as practicing Christians and Muslims respectively. Only 3.4% indicated that they had no religion. Twenty-three participants (19.8%) had paid employment in addition to their studentships. Most of the participants reported that, they were fully supported financially by their relatives (42.2%). Others reported that, they were supported by their parents (19.8%), friends (5.2%) and government scholarship (18.1%). The rest (14.7%) indicated that they were self-sponsored.

**Table 1:** Socio-demographic Characteristics of Participants (*n* = *116*).

| Variable | Number | % or Mean ± SD |
| --- | --- | --- |
| **Gender** |  |  |
| Male | 75 | 64.70% |
| Female | 41 | 35.30% |
| **Age (years)** |  |  |
| All | 116 | $24.6 \pm 5.7$ |
| Male | 75 | $25.7 \pm 5.7$ |
| Female | 41 | $22.7 \pm 5.1$ |
| **Marital Status** |  |  |
| Married | 20 | 17.20% |
| Single | 52 | 44.80% |
| Divorced | 2 | 1.70% |
| In a Relationship | 42 | 36.20% |
| **Occupation** |  |  |
| Student Only | 93 | 80.20% |
| Part-time Job | 23 | 19.80% |
| **Practice of Religion** |  |  |
| Christian | 93 | 80.20% |
| Muslim | 19 | 16.40% |
| Others | 4 | 3.40% |
| **Residence** |  |  |
| University Hostel | 12 | 10.30% |
| Renting a room/house | 95 | 81.90% |
| Living with Parent/Relatives | 9 | 7.80% |
| **Financial Support** |  |  |
| Parents | 23 | 19.80% |
| Friends | 6 | 5.20% |
| Relatives | 49 | 42.20% |
| Government Scholarship | 21 | 18.10% |
| Self-sponsored | 17 | 14.70% |
| **Nationality** |  |  |
| Uganda | 91 | 78.50% |
| Kenya | 12 | 10.30% |
| Nigeria | 12 | 10.30% |
| Rwanda | 1 | 0.90% |
| **Family Type** |  |  |
| Monogamous | 81 | 69.90% |
| Polygamous | 35 | 30.10% |
| **Quality of Relation with Parents** |  |  |
| Good | 94 | 81.00% |
| Moderate | 19 | 16.40% |
| Poor | 3 | 2.60% |
| **Parental loss** |  |  |
| Both are Alive | 48 | 41.10% |
| One Parent is Alive | 26 | 22.40% |
| Orphan | 42 | 36.20% |
| **Age at Parental Loss** |  |  |
| ≤ 10 years | 12 | 17.60% |
| > 10 years | 56 | 82.40% |

### Prevalence of depressive symptom among participants

Table 2 shows the proportion of participants whose BDI score indicated depressive symptoms. The results indicated that 33 (28.4%) of the 116 participants exhibited depressive symptoms. According to the cut off scores, those that were detected as having depressive symptoms consisted of 16 (13.8%) cases of mild mood disturbance (BDI score of 10-18), 14 (12.1%) cases of moderate depression (BDI score of 19-29) and 3 (2.6%) cases of severe depression (BDI score of 30-40). There was no reported case of very severe depression (BDI score of >40).

**Table 2:** Percentages of participants whose BDI Score indicated depression by gender (Mean = 9.04, SD = 7.4, Range: 1 – 45)

| BDI score | Male (n = 75) | Female (n = 41) | Total (n = 116) |
| --- | --- | --- | --- |
| **Normal (0-9)** | 56 (48.3%) | 27 (23.2%) | 83 (71.6%) |
| **Mild Mood Disturbance (10 -18)** | 10 (8.6%) | 6 (5.2%) | 16 (13.8%) |
| **Moderate Depression (19-29)** | 8 (6.9%) | 6 (5.2%) | 14 (12.1%) |
| **Severe Depression (30- 40)** | 1 (0.9%) | 2 (1.7%) | 3 (2.6%) |

The incidence of depression was found to be more among male participants (16.4%) when compared to female participants (12.1%) as shown in Fig 1. There were no significant differences in mean depression scores in relation to age, sex, marital status, financial support or type of residence (p >0.05). Among the depressed participants, 9 (7.8%) reported having suicidal ideation over the past 2 weeks as compared to 8 (6.8%) of participants without depressive symptoms (Table 3).

**Figure 1:**
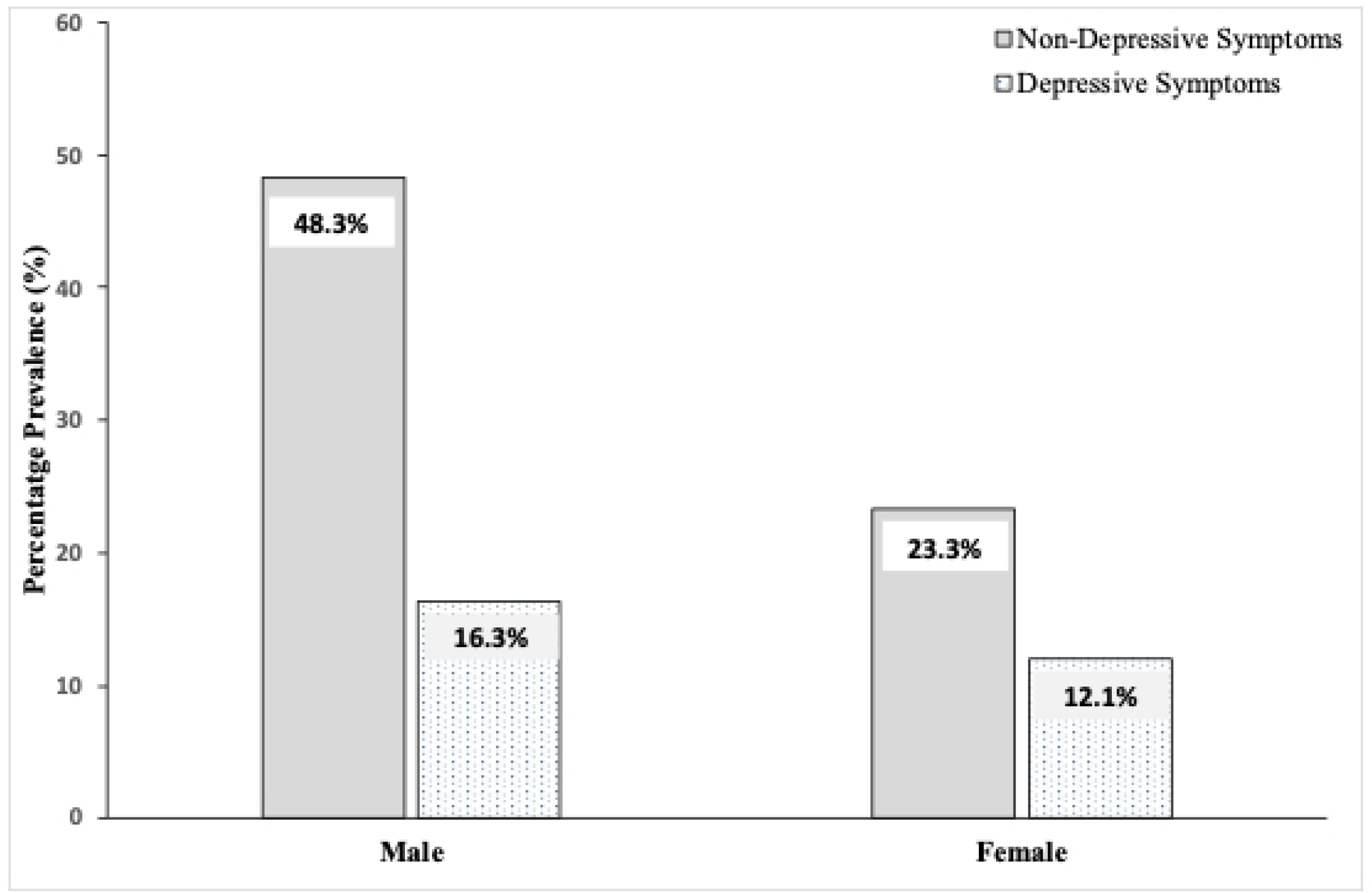
Prevalence of Depression among Participants (*n* = *116*)

**Table 3:** Bivariate Analysis of Risk Factors for Depressive Symptoms

| Variable | Depressive Symptoms | Non-Depressive Symptoms | X<sup>2</sup> (df) | P-value |
| --- | --- | --- | --- | --- |
|  | n (%) | n (%) |  |  |
| Suicidal Ideation (n=116) |  |  |  |  |
| Yes | 9 (7.6) | 8 (6.8) | 17.005 (1) | 0.0003 |
| No | 24 (20.7) | 75 (64.5) |  |  |
| Age Group in years (n =116) |  |  |  |  |
| < 20 | 3 (2.6) | 5 (4.3) | 1.974 (3) | 0.578 |
| 20 - 29 | 25 (21.6) | 63 (54.3) |  |  |
| 30 - 39 | 3 (2.6) | 13 (11.2) |  |  |
| 40 - 49 | 2 (1.7) | 2 (1.7) |  |  |
| Gender (n=116) |  |  |  |  |
| Male | 19 (16.4) | 56 (48.3) | 1.011 (1) | 0.315 |
| Female | 14 (12.1) | 27 (23.3) |  |  |
| Residence (n=116) |  |  |  |  |
| University Hostel | 5 (4.3) | 7 (6) | 1.376 (2) | 0.307 |
| Renting a room/house | 27 (23.3) | 68 (58.6) |  |  |
| Living with Parents or Relatives | 1 (0.9) | 8 (6.9) |  |  |
| Financial Support (n=116) |  |  |  |  |
| Friends | 2 (1.7) | 4 (3.4) | 1.818 (4) | 0.769 |
| Government Scholarship | 8 (6) | 13 (12.1) |  |  |
| Parents | 3 (3.4) | 20 (16.4) |  |  |
| Relatives | 15 (12.9) | 34 (29.3) |  |  |
| Self-sponsored | 5 (4.3) | 12 (10.3) |  |  |
| Marital Status (n=116) |  |  |  |  |
| Single | 16 (13.8) | 36 (31.0) | 7.418 (3) | 0.058 |
| In a relationship | 9 (7.8) | 33 (28.4) |  |  |
| Married | 6 (5.1) | 14 (12.3) |  |  |
| Divorced | 2 (1.7) | 0 (0) |  |  |
| Occupation (n=116) |  |  |  |  |
| Student only | 24 (20.7) | 69 (59.5) | 1.608 (1) | 0.205 |
| Part-time job | 9 (7.8) | 14 (12.1) |  |  |
| Family Type (n=116) |  |  |  |  |
| Monogamous | 22 (19) | 59 (50.9) | 0.219 (1) | 0.640 |
| Polygamous | 11 (9.5) | 24 (20.7) |  |  |
| Parent Loss (n=116) |  |  |  |  |
| Both are alive | 12 (10.3) | 36 (31.0) | 0.905 (2) | 0.636 |
| Only one is alive | 7 (6.0) | 19 (16.4) |  |  |
| Orphan | 14 (12.1) | 28 (24.1) |  |  |
| Quality of Relationship with Parents (n=116) |  |  |  |  |
| Good | 23 (19.8) | 71 (61.2) | 7.042 (2) | 0.0296 |
| Moderate | 9 (7.6) | 10 (8.6) |  |  |
| Poor | 1 (0.9) | 2 (1.7) |  |  |

### Relationship between depression and associated risk factors

The association between BDI scores with different psychosocial and demographic variables was determined by bivariate and multivariate analysis. The results obtained for the bivariate analysis are displayed in Table 3. The analysis did not indicate significant differences in participants with depressive symptoms when compared to those without symptoms of depression in relation to gender (*X*^2^ = 1.011, df = 1, p = 0.315), age (*X*^2^ = 1.974, df = 3, p = 0.578), type of residence (*X*^2^ = 1.376, df = 2, p = 0.307), type of financial support (*X*^2^ = 1.818, df = 4, p = 0.769) and family type (*X*^2^ = 0.219, df = 1, p = 0.640). However, there was a weak and significant association of depressive symptoms with marital status (*X*^2^ = 7.418, df = 3, p = 0.058) and the thoughts of committing suicide (*X*^2^ = 17.005, df = 1, p = 0.0003); and relationship with parents (*X*^2^ = 7.042, df = 2, p = 0.0296) respectively.

Results from multivariate analysis indicates that, the type of residence where the participant is staying in (OR 1.94, 95 % CI 0.57 – 6.61, p < 0.05), the thoughts of committing suicide (OR 15.19, 95 % CI 3.07 – 75.11, p < 0.05), and the source of financial support the individual is receiving (OR 1.33, 95 % CI 0.48 – 3.65, p < 0.05) were each independently associated with significant depressive symptoms.

### Mean levels of 24-hour urinary norepinephrine excretion

The mean 24-h NE excretion levels of all the depressed participants (121.97 ± 51.48 μg/day) was higher when compared to participants without depressive symptoms (87.58 ± 18.64 μg/day) but the difference was not statistically significant (P > 0.05, Fig 2). 24-h urinary NE concentrations in females and males were not remarkably different in participants with or without depressive symptoms (p > 0.05). However, 29 of the 33 participants with depressive symptoms showed NE levels above the reference range. In addition, the mean amount of NE excreted by participants with depressive symptoms was not related with the severity of depression (MMD = 118.13±38.99 μg/day, MD = 124.79±36.48 μg/day, SD = 129.33±39.56 μg/day) respectively (Fig 3).

**Figure 2:**
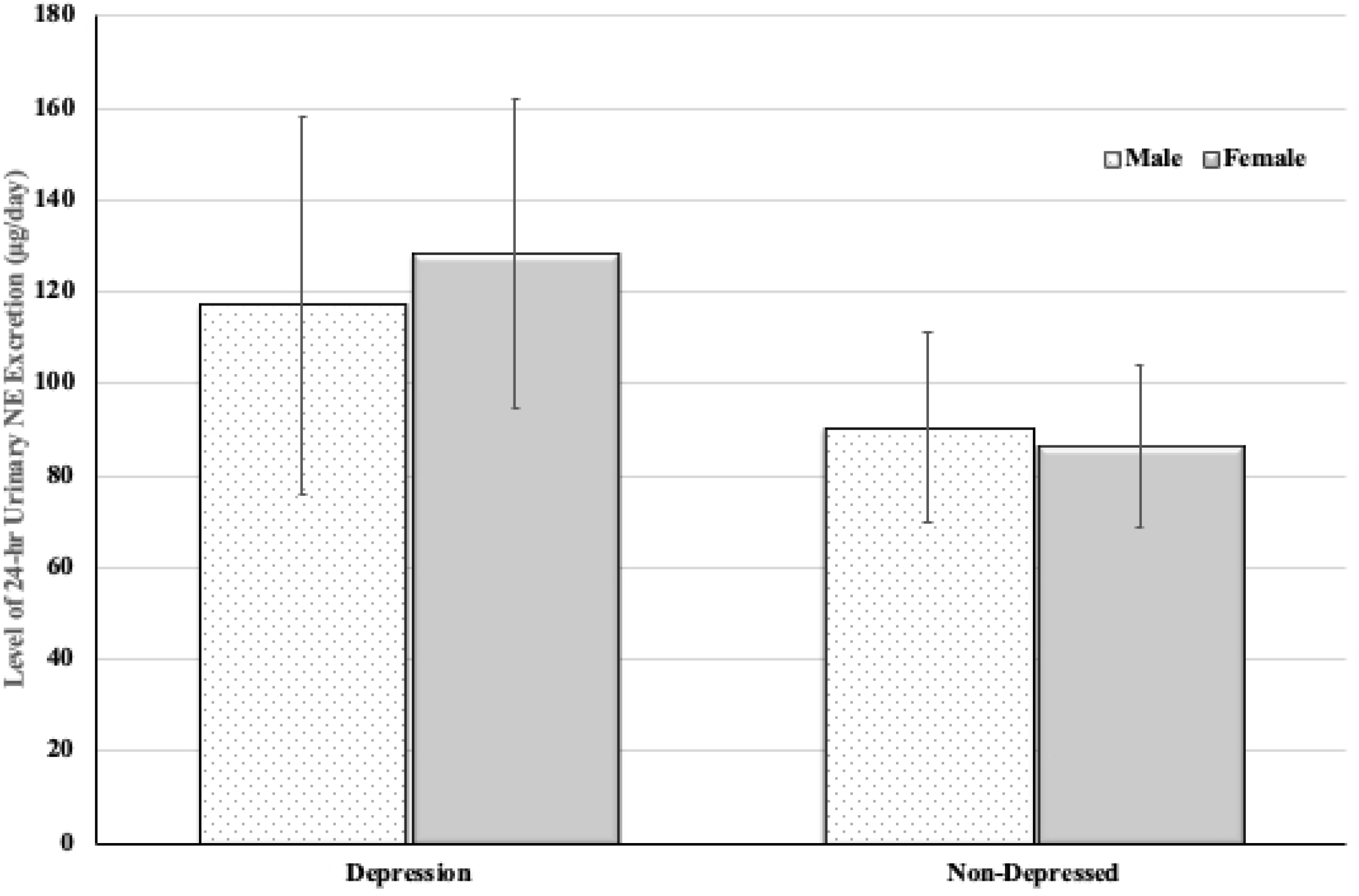
Mean (± SD) 24 – h of urinary excretion (μg/ml) of norepinephrine of the study population (n= 116).

**Figure 3:**
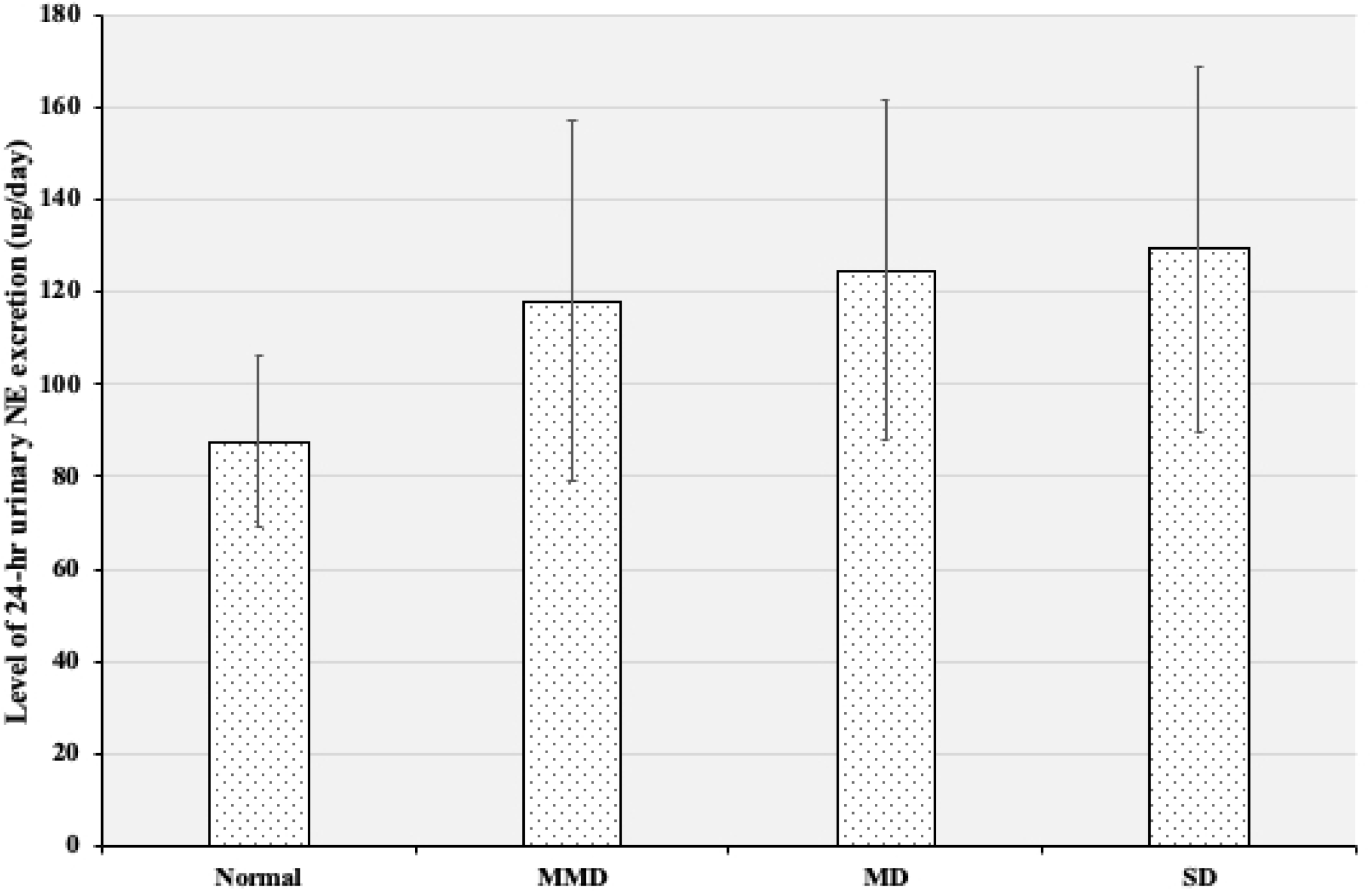
Mean (± SD) 24 – h of urinary excretion (μg/ml) of norepinephrine of the study population according to Beck Depression Inventory (BDI) score (n= 116). *Mild Mood Disturbance (MMD)*, *Moderate Disturbance (MD)*, *Severe Depression (SD)*

A simple linear regression analysis was done to determine the relationship between the level of NE and the occurrence of depressive symptomatology. The results showed that there was a weak non-significant correlation between the BDI and the 24-h urinary NE level [coefficient of correlation (r^2^ = 0.2059, P > 0.05; Fig 4). In the calculated relative risk analysis, males were 1.2 (95% CI 0.612 – 2.423) times more likely to have higher urinary NE excretion levels compared to females.

**Figure 4:**
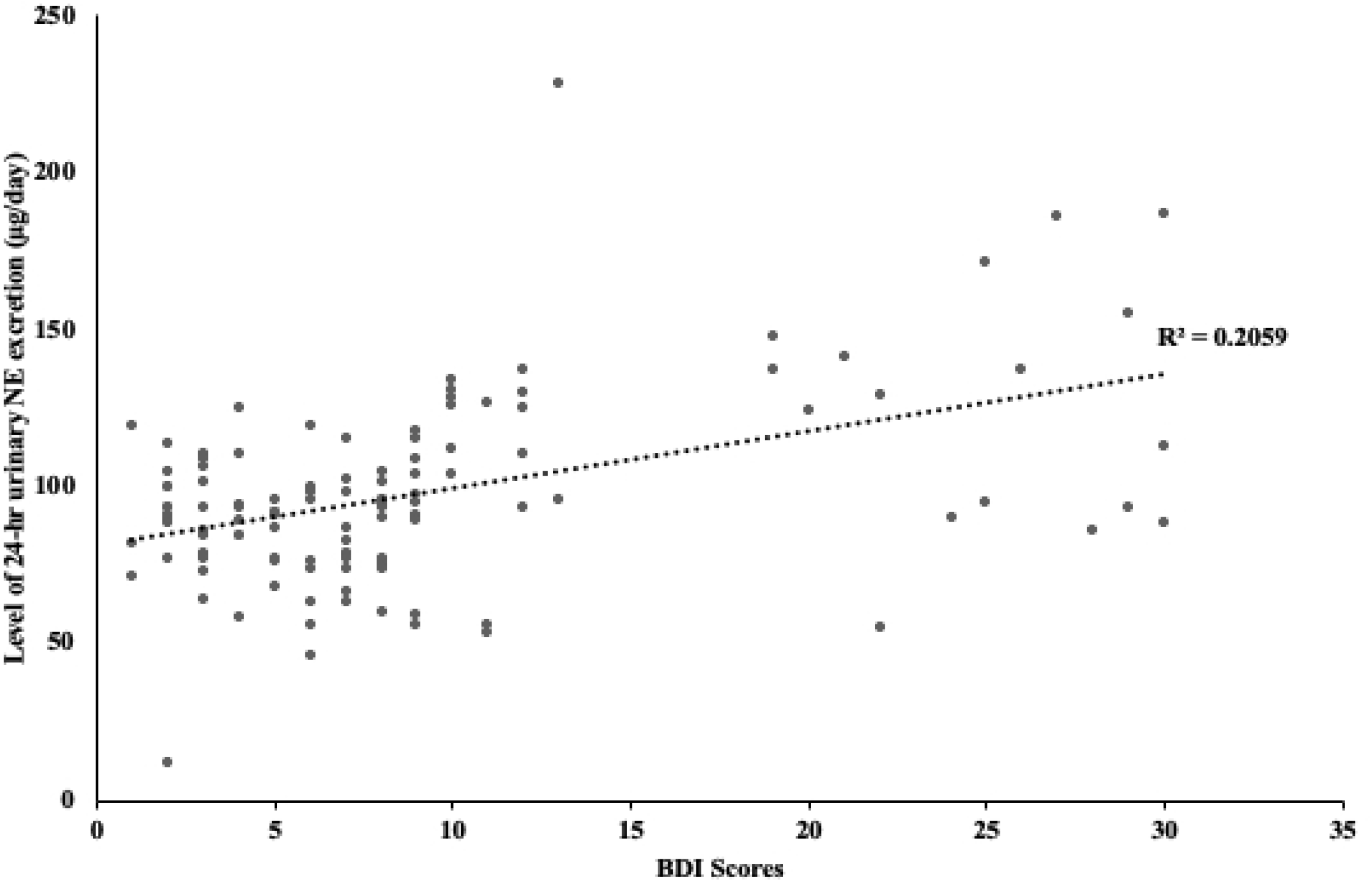
Linear regression coefficient of the relation between 24-h urine norepinephrine (NE) concentrations and the presence of depressive symptoms.

## Discussion

To the best of our knowledge, this is the first study to estimate the prevalence of depression and examine the association with socio-demographic and biological determinants of the disease among undergraduate medical students in Uganda. Only one previous study has determined the prevalence of depression among university students in Uganda without examining the underlying cause of the observed rate of depression [12]. The results of our study indicated that, 28.4% and 14.4% of undergraduate medical students of Kampala International University, Uganda, exhibited depressive symptoms and suicide ideation respectively. No significant changes in mean depression scores were observed among male and female participants even though prevalence was slightly higher in males. The participants with depressive symptoms were more likely than those without symptoms to have increased levels of 24-h mean NE excretion above the normal range (<90 μg/day).

The rate of depression of 28.4% is much higher than the 16.24% reported earlier among the first-year students of Makerere University Kampala, Uganda [12] but closely aligned with the results obtained from meta-analysis of 195 studies involving 129,123 medical students in 47 countries which demonstrated that 27.2% of medical students (range, 9.3% – 55.9%) screened positive for depression and that 11.1% (range, 7.4% – 24.2%) reported suicidal ideation [31]. The prevalence of depression and percentage of individuals with suicide ideation observed in this study is concerning given that the development of depression and suicidality has been linked to an increased short-term risk of suicide as well as a higher long-term risk of future depressive episodes and morbidity [32, 33]. Also, depressive and suicidal symptoms in medical trainees may adversely affect the long-term health of physicians as well as the quality of care [34].

The concept that depression can be caused only by psychosocial or biological factors have been suggested but their causal links still remain unclear. We found that depression rates were higher among students within the age group of 20 – 29 years (21.6%), those renting a room/house outside the university (23.3%), those who are full-time students (20.7%), those whose financial support comes from relatives (12.9%), and surprisingly those who claimed to have good quality relationship with parents (21.6%) when compare to other variables in the same category (Table 3). When these variables were examined as risk factors for depression in the bivariate analysis, marital status, relationship with parents and suicide ideation prove significant. In Multivariate analysis, the type of residence, source of financial support and suicide ideation were each independently associated with significant depressive symptoms. We observed no interactions between depressive symptoms and the other variables that were entered in our logistic model (all p values for interaction > 0.05). Similar studies elsewhere indicated problems linked with accommodation, very large family size, financial hardships, difficulties in relationships, heavy cigarette smoking and high level of alcohol consumption, and fear of examinations were significantly associated with depressive disorders [35–37]. Although studies involving similarly aged members in the general population have reported a higher prevalence of depression among women compared to men [38,39], we found no evidence that women were more likely than men to experience depression as our results indicated that depressive symptoms were slightly higher in males than females. This is similar to the observations of Bostanci et al., and Bayram et al., [40, 41] who reported no difference between depression and gender.

While urinary measures are not a direct assessment of central activity, studies have characterized urinary neurotransmitters as biomarkers of various conditions linked to the disruptions within the central nervous system [25, 42]. The causal direction between BDI scores and NE levels was assessed in this study. The results showed a positive non-significant correlation between the BDI scores and 24-h urinary NE level (r^2^ = 0.2059, P > 0.05). In stepwise multiple linear regression tests, 24-h urinary NE concentration was not influenced by age or any of our predetermined associated risk factors. Other studies have consistently show significant relationship of depression disorders with urinary NE excretion levels [43, 44). The observed elevated NE levels may be associated with increased HPA-axis activity as previously reported [45]. An increase in hypothalamic–pituitary–adrenal (HPA) activity has been observed in 20% to 40% of depressed outpatients and in 40% to 60% of depressed inpatients [46]. Since all the participants in the present study were not suffering from any diseases associated with the cardiovascular or the endocrine systems, the elevated NE levels in the integrated 24-h urine samples indicates a plausible causative role of NE in increasing the risk of developing depressive symptoms. These findings argue for assessment of urinary NE excretion in the diagnosis of depressive disorders in the context of a detailed patient history.

From the foregoing, the findings of this study indicate that biological and psychosocial factors may be considered as risk factors within a larger framework for explaining the etiology of depression. The study is limited by its cross-sectional nature and the fact that only one medical school out of the five in the country at the time of the study was represented. In addition, the study variables were measured by self-report questionnaires, which do not allow diagnostic conclusions since there was no secondary screening such as a clinical interview. However, these self-reported inventories are essential tools for accurately measuring depression because they protect anonymity in a manner that is not possible through formal diagnostic interviews [47]. Due to budget constrain, this study was only able to assess the biomedical student population neglecting students in the clinical years. As a result, there was no comparison with students of other academic years in the University or with another University in different part of the country. The strength of our study was in it being the first to examine the prevalence of depression and 24-h urinary NE level among university students in Uganda.

## Conclusions

In conclusion, we found that the prevalence of depression or depressive symptoms among Kampala International University medical students was 28.4% which was within the rates reported from other countries. Increased levels of NE excretion and psychosocial factors may contribute to an increased risk of developing depressive symptoms in the population studied. Because of the high prevalence of depressive and suicidal symptomatology observed among medical students, it is important for medical schools to recognize and support all students experiencing depression, but in particular to consider how best to encourage this group to seek help. Further research is needed to identify strategies for preventing, identify causes of emotional distress and treating these disorders.

## Acknowledgement

The authors would like to sincerely thank the participants in the study who took time to complete our questionnaire and sacrifice their weekends to provide urine despite our stringent research protocol. Also special thanks to Dr. Martha Crespo Vicente for her advice and making available the laboratory space and equipment in the Institute of Biomedical Research, Kampala International University, Uganda.

## Supporting information

S1 Fig 1.

S2 Fig 2.

S3 Fig 3.

S4 Fig 4.

S5 Table 1.

S6 Table 2.

S7 Table 3.

## Data availability

All relevant data are within the paper.

